# Mutations in NBEAL2 do not impact Weibel-Palade body biogenesis and Von Willebrand factor secretion in Gray Platelet Syndrome Endothelial Colony Forming Cells

**DOI:** 10.1101/2022.06.08.495181

**Authors:** Marije Kat, Iris van Moort, Petra E. Bürgisser, Taco W. Kuijpers, Menno Hofman, Marie Favier, Rémi Favier, Coert Margadant, Jan Voorberg, Ruben Bierings

## Abstract

**Background:** Gray Platelet Syndrome (GPS) patients with Neurobeachin-like 2 (NBEAL2) deficiency produce platelets lacking alpha-granules (AGs) and present with lifelong bleeding symptoms. AGs are lysosome-related organelles (LROs) and store the hemostatic protein Von Willebrand factor (VWF) and the transmembrane protein P-selectin. Weibel-Palade bodies (WPBs) are LROs of endothelial cells and also store VWF and P-selectin. In megakaryocytes, NBEAL2 links P-selectin on AGs to the SNARE protein SEC22B on the endoplasmic reticulum (ER), thereby preventing premature release of cargo from AG precursors. In endothelial cells, SEC22B drives VWF trafficking from ER to Golgi and promotes the formation of elongated WPBs, but it is unclear if this requires NBEAL2.

**Objectives:** To investigate a potential role for NBEAL2 in WPB biogenesis and VWF secretion using NBEAL2 deficient endothelial cells.

**Methods:** Interaction of SEC22B with NBEAL2 in endothelial cells was investigated by interactomic mass spectrometry and pull down analysis. Endothelial Colony Forming Cells (ECFCs) were isolated from healthy controls and 3 unrelated GPS patients with mutations in *NBEAL2*.

**Results:** We show that SEC22B binds to NBEAL2 in ECs. GPS patient-derived ECFCs are deficient of NBEAL2, but reveal normal formation and maturation of WPBs and normal WPB cargo recruitment. Neither basal nor histamine-induced VWF secretion are altered in the absence of NBEAL2.

**Conclusions:** While NBEAL2 deficiency causes absence of AGs in GPS patients, it has no impact on WPB functionality in ECs. Our data highlight the difference in regulatory mechanisms between these two hemostatic storage compartments.

**Essentials:**

1. We characterized Gray Platelet Syndrome patient-derived endothelial cells with biallelic NBEAL2 mutations *ex vivo*.
2. NBEAL2 is not essential for Weibel-Palade body biogenesis, maturation, and Von
Willebrand factor secretion from endothelial cells.

## Introduction

Gray platelet syndrome (GPS) is a rare platelet storage pool disorder (SPD) that is characterized by thrombocytopenia, enlarged platelets and lack of platelet alpha-granules (AGs), causing increased bleeding tendency, myelofibrosis and splenomegaly.^1^ AGs store bioactive compounds, including Von Willebrand Factor (VWF), a crucial protein for hemostasis and wound healing.^2^ Weibel-Palade bodies (WPBs) are the endothelial cell (EC) equivalent of AGs and contain VWF as their main cargo.^3^ Both AGs and WPBs are lysosome-related organelles (LROs), a family of cell-type specific (secretory) organelles that acquire cargo and membrane components from the endolysosomal system and which have synergistic functions in immunity, hemostasis, inflammation and angiogenesis.^4,5^ LRO biogenesis and degranulation are in many respects regulated by universal mechanisms that are shared between cell types that contain LROs.^4^ Genetic defects causing SPDs often affect LRO biogenesis and release in multiple cell types simultaneously, including endothelial WPBs.^5^ Vascular injury triggers VWF secretion from WPBs, producing adhesive strings that mediate platelet adherence to the damaged vessel wall.^3^ Platelets subsequently become activated and release AG content to stabilize the platelet plug.^5^ Evidently, in GPS AG degranulation is absent, resulting in platelet aggregation defects and bleeding complications in patients.

GPS is caused by autosomal recessive mutations in neurobeachin-like 2 (NBEAL2),^6–8^ while autosomal dominant mutations in the transcription factor Growth Factor Independent 1B (GFI1B) underlie a related ‘GPS-like’ syndrome.^9^ NBEAL2 is required for AG biogenesis, potentially through retention of cargo proteins in AG precursors during maturation or by retention of AGs flowing their formation in megakaryocytes.^10,11^ GPS neutrophils also exhibit defects in a subset of their LROs, the specific granules (SGs), which include loss of SG cargo due to premature release of SGs.^12^ Taken together this suggests that NBEAL2 potentially serves a general role in regulation of LROs.^12^ The SNARE protein SEC22B is a central player in transport between the endoplasmic reticulum (ER) and the Golgi ^13^ that has recently been implicated in AG and WPB biogenesis. We have previously uncovered that in ECs, SEC22B controls the formation and morphology of WPBs.^14,15^ In megakaryocytes, NBEAL2 links SEC22B to the leukocyte receptor P-selectin present on AGs in a tripartite complex which acts to prevent the premature release of AG cargo from AG precursors through a mechanism that needs further clarification.^16^ Given the prominent role of SEC22B in WPB biogenesis and because P-selectin is also present on WPBs^17^, we investigated whether NBEAL2 plays a similar role in the endothelial secretory pathway.

## Materials and Methods

### Cell culture and isolation of endothelial cells

Pooled, cryopreserved human umbilical vein endothelial cells (HUVECs) were purchased from Promocell. Endothelial colony forming cells (ECFCs) were isolated from heparinized venous blood as described.^18^ Control ECFCs were isolated from healthy, pseudonymized volunteers participating in Sanquin’s internal blood donor system. GPS ECFCs were isolated from three unrelated GPS patients, coded C (6 clones), D (2 clones) and E (2 clones), that were previously described ^12^, and correspond to patient C, D and E from that study. The study was performed according to national regulations regarding the use of human materials, informed consent and the Declaration of Helsinki and was approved by the Medical Ethical Committee of the Academic Medical Center in Amsterdam. HUVECs and ECFCs were cultured as described.^19^ Expression of interleukin-8 (IL-8) was upregulated by incubation with 10 ng/ml interleukin-1β (IL-1β) as described previously.^20^ Immunocytochemistry, immunoblotting and VWF secretion assays were performed essentially as described.^19,21^ Antibodies and dilutions are described in the Supplemental Materials and Methods.

## Results and Discussion

### NBEAL2 interacts with SEC22B in primary ECs

NBEAL2 was previously shown to be part of a tripartite complex linking SEC22B and P-selectin in megakaryocytes.^16^ We have recently performed an interactomic analysis of SEC22B in ECs, which identified the SNARE protein STX5 as a novel regulator of WPB size as well as VWF trafficking and secretion.^22^ We revisited that dataset, which is publically accessible via the PRIDE repository with dataset identifier PXD027516, and found that NBEAL2 was also identified as a specific interactor of SEC22B in ECs (**Figure 1A**). To confirm this we used mEGFP-SEC22B as bait in a pulldown assay in HUVECs followed by immunoblot analysis. Because ECs also contain P-selectin and, like megakaryocytes, express it on the limiting membrane of their LROs (i.e. WPBs), we also investigated whether the tripartite SEC22B/NBEAL2/P-selectin complex is present in ECs. We indeed confirmed a specific interaction with endogenous NBEAL2, but found no evidence of such a tripartite complex in ECs as P-selectin coprecipitation was not observed (**Figure 1B**).

**Figure 1.**
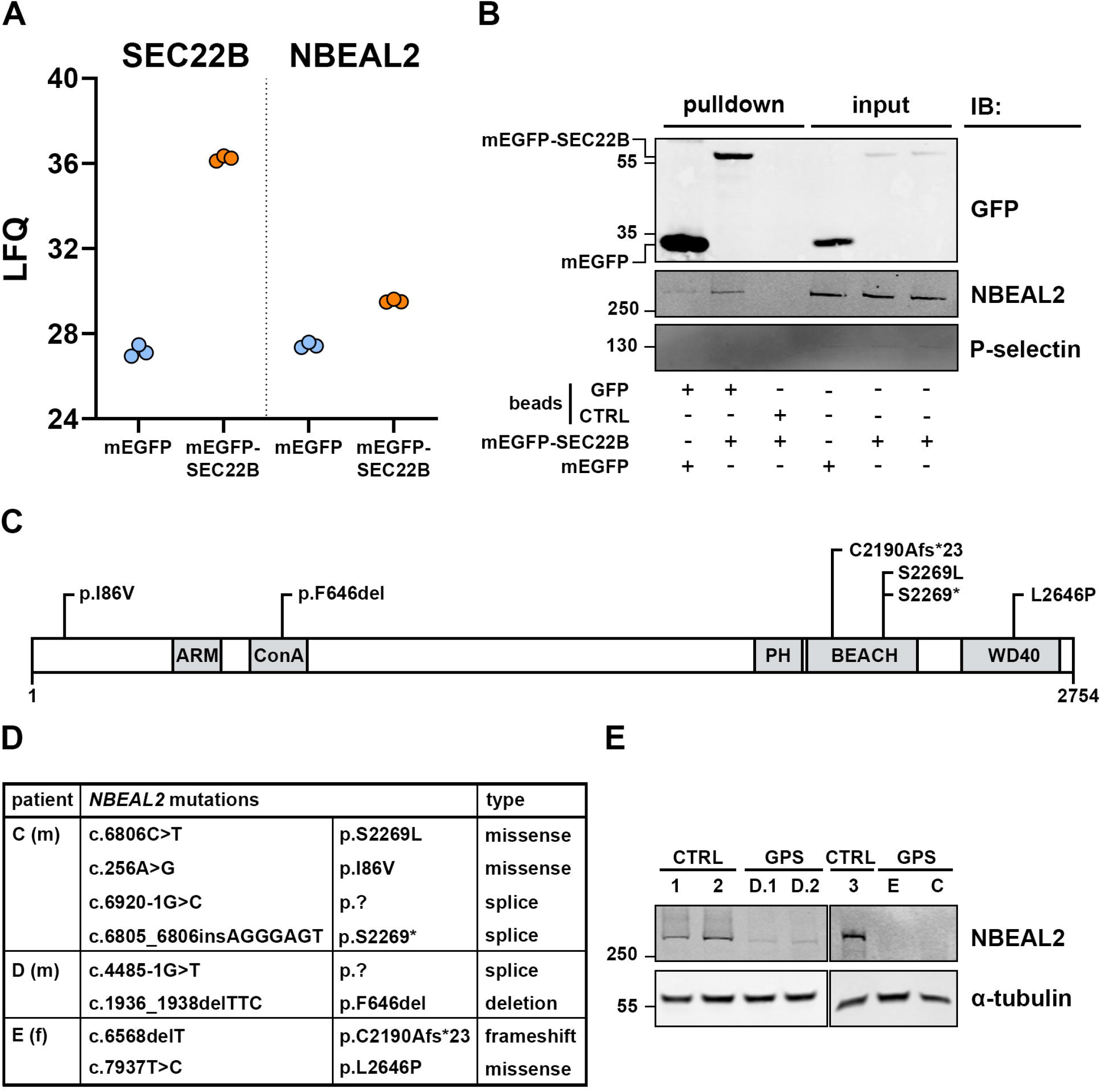
Mutations in the SEC22B interactor NBEAL2 are associated with loss of NBEAL2 expression in GPS patient-derived endothelial cells. (A) Label-free quantification (LFQ) of SEC22B and NBEAL2 proteins levels in GFP pull down samples of mEGFP or mEGFP-SEC22B baits expressed in HUVECs determined by mass spectrometry analysis.^22^ (B) Pull down analysis of mEGFP or mEGFP-SEC22B bait expressed in HUVECs using GFP or control (ctrl) beads and Western blot analysis of GFP, NBEAL2, and P-selectin. Molecular weights of protein ladder marker bands are indicated in kDa on the left. (C) Domain structure of full length (2754 a.a.) NBEAL2 showing the position of the mutations present in patients C, D and E. (D) Table showing *NBEAL2* gene mutations, changes on protein level and type of mutation in GPS patients included in this study (m: male, f: female). Gene transcript identifier: NM_015175 / Protein identifier: NP_055990. (E) Western blot of NBEAL2 expression in ECFCs from 3 GPS patients (C, D and E; 2 clones were analyzed for patient D) compared to ECFCs isolated from healthy control donors (CTRL 1-3). α-tubulin is shown as a loading control. Molecular weights of protein ladder marker bands are indicated in kDa on the left.

### NBEAL2 deficient ECFCs produce WPBs

A previous study showed that ECs in tissue sections of patients lacking platelet AGs have WPBs containing VWF and P-selectin.^23^ However, it has since been shown that defects in NBEAL2 and GFI1B both lead to AG loss,^6,7,24^ and it was unclear whether these patients were NBEAL2 deficient. To investigate the role of NBEAL2 in WPB regulation, we isolated ECFCs from three GPS patients with different mutations in *NBEAL2* (**Figure 1C,D**). These patients were previously shown to have platelets with no AGs, diminished AG cargo and elevated steady-state cell surface P-selectin ^12,25^, and also neutrophils with reduced SG cargo (including lactoferrin), elevated steady-state surface expression of the SG protein CD66b and defective NETosis ^12^, which is thought to be the result of impaired AG and SG retention in platelets and neutrophils respectively. We first confirmed that GPS patient-derived ECFCs are NBEAL2-deficient using immunoblot analysis of ECFC lysates (**Figure 1E, Supplemental Figure 1**), which showed that NBEAL2 was undetectable in ECFCs of patient C and E and all but lost in ECFCs of patient D. Next we assessed the effect of *NBEAL2* mutations on WPB formation in cultured GPS ECFCs. Immunofluorescent staining of VWF revealed rod-shaped WPBs in all clones, with no apparent changes in number or morphology compared to those in healthy donor ECFCs (**Figure 2A**). Furthermore, in CTRL and GPS ECFCs SEC22B staining was observed on the ER and Golgi, while P-selectin was present on WPBs of CTRL as well as GPS ECFCs, suggesting that their intracellular localization is independent of NBEAL2 (**Figure 2B**). The lack of overlap between SEC22B and P-selectin further highlights that the presence of a tripartite SEC22B/NBEAL2/P-selectin complex in ECs is unlikely because of significant spatial separation. Immunolocalization of NBEAL2 resulted in a dispersed intracellular staining pattern with no overlap with VWF or resemblance to any other intracellular structures (**Supplemental Figure 2**).

**Figure 2.**
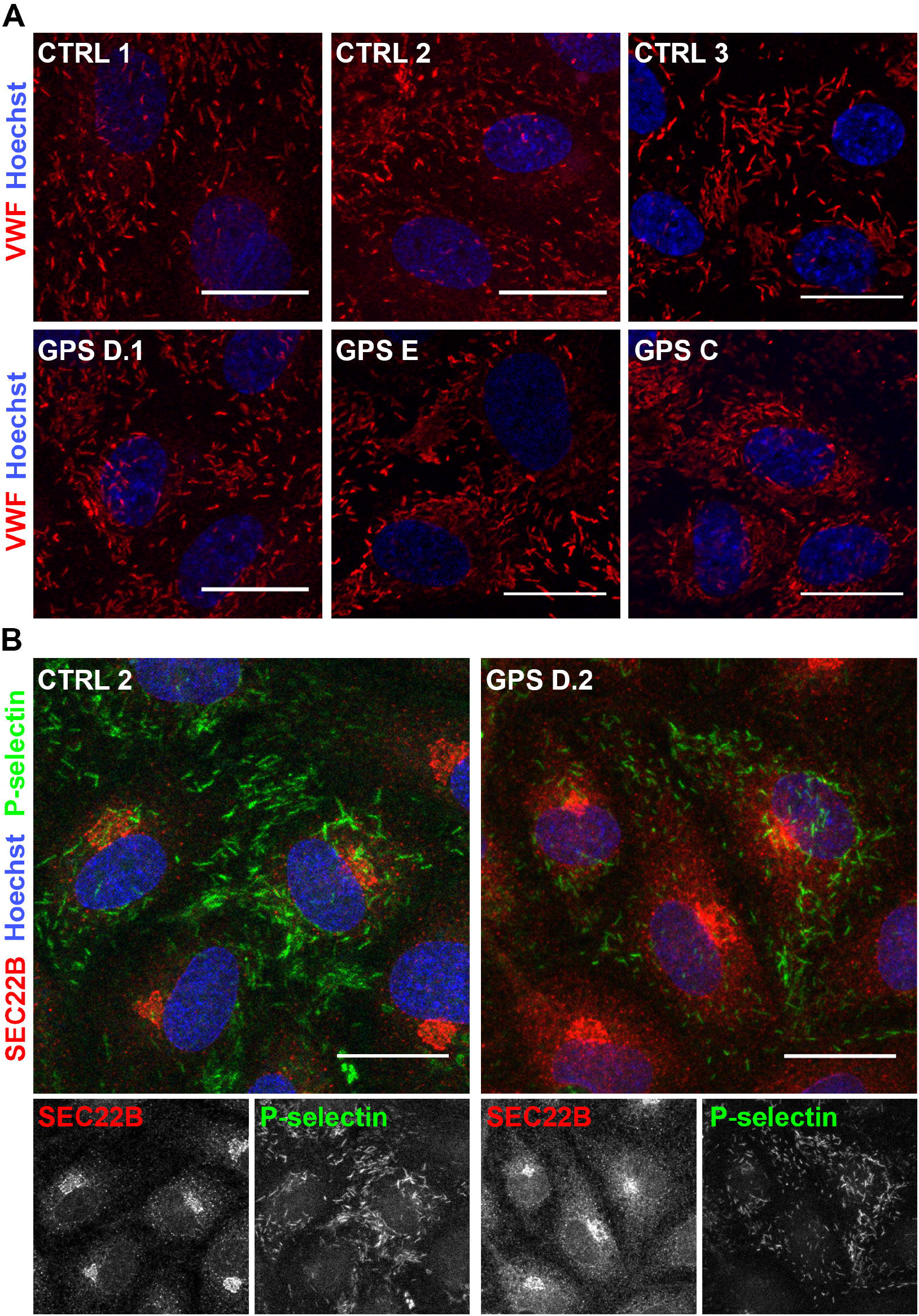
GPS patient-derived endothelial cells generate WPBs despite NBEAL2 deficiency. Representative maximal projections of confocal immunofluorescent analysis of (A) VWF (red) and nuclei (Hoechst, blue) and (B) SEC22B (red), P-selectin (green) and nuclei (blue) in CTRL and GPS patient-derived ECFCs. Grayscale images for SEC22B and P-selectin channels are shown below. Scale bars represent 20 μm.

Since a previous report suggested that endogenously synthesized (VWF) as well as endocytosed (fibrinogen) AG cargo proteins were not retained in AG precursors during AG biogenesis in NBEAL2 deficient megakaryocytes ^10^, we next tested whether the composition of WPBs was altered in GPS ECFCs. For this we examined the presence of 3 WPB cargo proteins: Angiopoietin 2 (Ang-2), which is endogenously synthesized and included into newly forming WPBs at the level of the *trans*-Golgi network (TGN), CD63, a tetraspanin that recycles from the plasma membrane and is delivered to WPBs via the endosomal compartment, and interleukin-8 (IL-8), a chemokine that is included in WPBs at the TGN after upregulation of its synthesis by interleukin-1β.^17,20,26–28^ IF stainings in CTRL and GPS ECFC clones revealed no change in Ang-2, CD63 and IL-8 localization to WPBs (**Figure 3A, Supplemental Figure 3A, Supplemental Figure 4**), which shows Golgi and post-Golgi WPB cargo recruitment and retention are unaffected by loss of NBEAL2. In addition, we observed normal recruitment of the small GTPase Rab27A, which is regarded as a maturation marker for WPBs and recruits effectors that act as positive and negative regulators of WPB exocytosis ^29^, indicating that this proximal event in recruitment of the WPB exocytotic machinery is not impaired by the absence of NBEAL2 (**Figure 3B, Supplemental Figure 3B**). Taken together our data suggest that NBEAL2 is not crucial for WPB biogenesis, cargo recruitment or maturation, hinting towards different regulatory mechanisms with regard to storage compartment biogenesis and (cargo) retention in comparison with megakaryocytes and platelets.^16^

**Figure 3.**
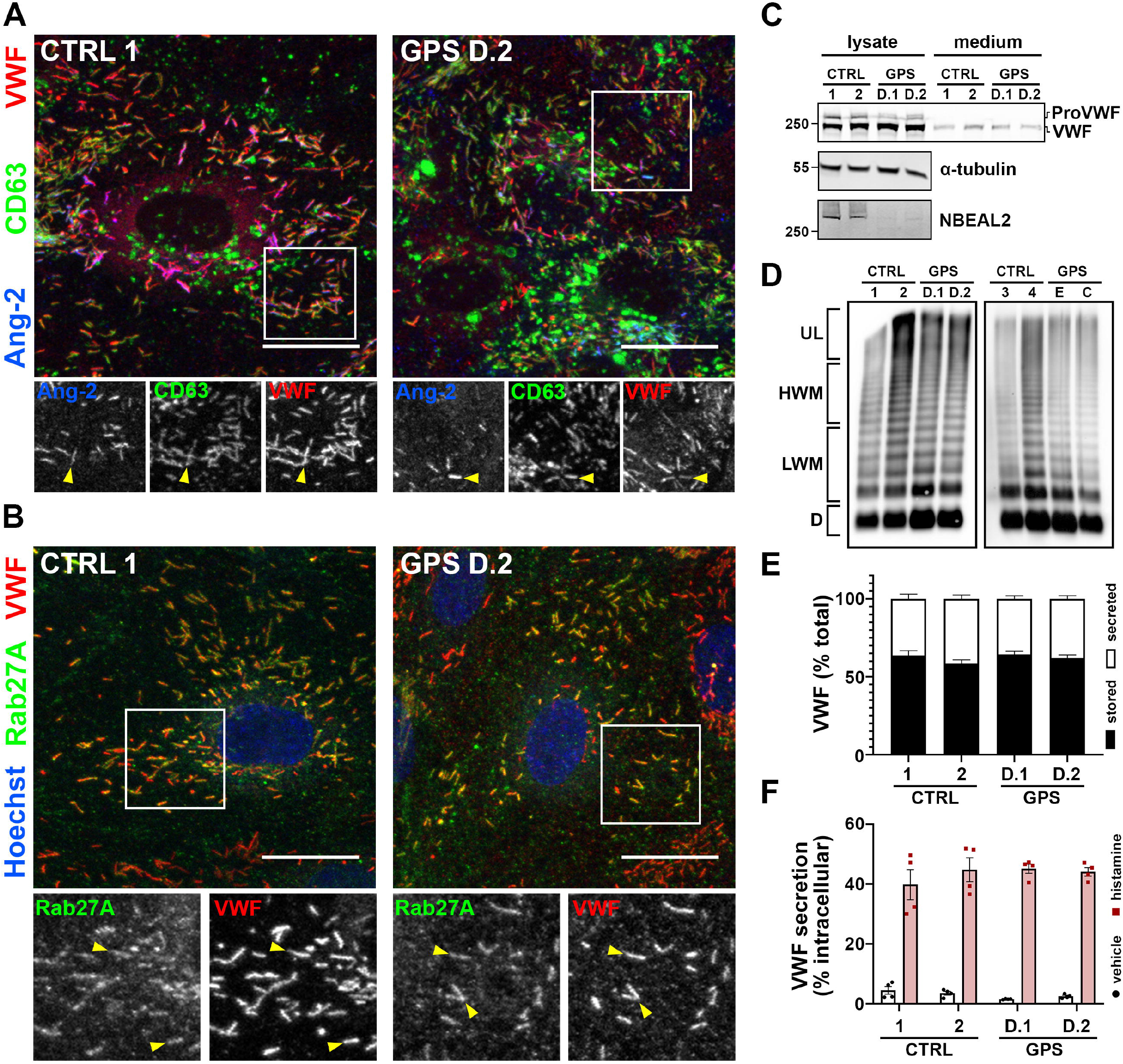
NBEAL2 is dispensable for WPB cargo sorting and maturation and for basal and secretagogue-induced VWF secretion. Representative maximal projections of confocal immunofluorescent analysis of (A) VWF (red), CD63 (green) and Angiopoietin 2 (Ang-2, blue) or (B) VWF (red), Rab27A (green), and nuclei (Hoechst, blue) in CTRL and GPS ECFCs. Boxed areas are shown below in grayscale. Yellow arrowheads indicate example WPBs. Scale bars represent 20 μm. (C) Western blot analysis of VWF content in lysates and media from CTRL and GPS patient-derived ECFCs. (D) VWF multimer analysis of 24 hour basal media from CTRL and and GPS patient-derived ECFCs. UL: ultra-large, HMW: high molecular weight, LMW: low molecular weight, D: dimer. (E) Percentage of stored and secreted VWF in 24 hours under basal conditions (n=7, mean±SEM, two-way ANOVA, not significant). (F) Secreted VWF in 30 minutes under unstimulated conditions (vehicle) or in the presence of 100 μM histamine expressed as a percentage of the total intracellular VWF levels in unstimulated cells (n=4, mean±SEM, two-way ANOVA, not significant).

### NBEAL2 is dispensable for basal and regulated VWF secretion

Since GPS is associated with a bleeding tendency^1^, we examined a possible contribution of disrupted endothelial VWF secretion. Accumulating evidence suggests that deficiency of platelet AGs and neutrophil SGs in GPS is underpinned by premature release of these granules or their cargo from megakaryocytes and neutrophils, respectively, due to defective retention.^10,12,25^ After biosynthesis WPBs are stored in ECs until exocytosis in response to activation, while at steady-state WPBs will stochastically undergo basal secretion.^3^ To test whether NBEAL2 deficiency impacts WPB retention at steady state or during stimulated secretion, we examined VWF storage and secretion during basal and histamine-induced release. No difference was seen in composition of intracellular or secreted VWF, with regard to proteolytic processing of ProVWF to mature VWF (**Figure 3C**) or to multimeric composition of secreted VWF (**Figure 3D**), indicating normal regulation of synthesis and secretion. Indeed, both basal (unstimulated) VWF secretion during 24 hours and 30-minute histamine-stimulated VWF secretion remained unchanged (**Figure 3E,F**). As VWF storage and secretion from NBEAL2 deficient ECs remains functional, this further points towards platelet defects as the prime cause for bleeding symptoms in GPS.^1^ Our findings that basal VWF release, which accounts for the bulk of circulating VWF, and histamine-stimulated VWF release are unaffected in GPS ECFCs are also consistent with the observations that GPS patients and *Nbeal2*^-/-^ mice have normal plasma levels of VWF ^30,31^ and that DDAVP infusion increases plasma VWF in GPS patients.^32^

Our results show that NBEAL2 is not required for WPB biogenesis nor WPB cargo retention, indicating that, contrary to its role in LRO cargo retention in megakaryocytes and neutrophils, in the endothelial secretory pathway it is nonessential. Tubular condensation of sufficient amounts of VWF in the lumen of the TGN is a key requirement for WPB biogenesis, and this shapes emerging WPBs into their characteristic elongated form.^15^ The essential role for VWF in this process is highlighted by the absence of WPBs in engineered and patient-derived VWF deficient endothelial cells ^26,33^ and the appearance of pseudo-WPBs after ectopic expression of VWF in non-endothelial cells.^34,35^ This dependence of WPB biogenesis on condensation of VWF is in contrast to the formation of platelet AGs, which are still present in platelets of patients with type 3 Von Willebrand Disease caused by mutations that abrogate VWF synthesis.^36,37^ We speculate that the difference between AG and WPB biogenesis in terms of NBEAL2 dependence may lie in the fact that WPBs are in many respects a nearly finished product after emergence from the TGN, while the cargo destined for AGs first proceeds through a number of endosomal compartments following its exit from the Golgi. During their synthesis in megakaryocytes, AGs are preceded by an intermediate stage that display the hallmarks of multi vesicular bodies (MVBs).^38^ These MVBs are late endosomal compartments that carry VWF originating from the TGN and will eventually progress to AG precursors, the site where NBEAL2 is thought to associate with P-selectin in megakaryocytes to retain AG cargo.^10^ Possibly, the molecular composition of the NBEAL2-dependent mechanism of LRO cargo retention differs between cell types, as is underscored by the notion that while neutrophils retain SGs and their cargo in a NBEAL2-dependent manner, they lack P-selectin.^12^ From a functional point of view, NBEAL2-dependent mechanisms that retain LROs or LRO cargo may be more crucial for cells such as neutrophils and megakaryocyte-derived platelets, which are unable to replenish secretory organelles after their initial synthesis, than for ECs that can continuously generate new WPBs.

## Acknowledgements

Work in our laboratory was funded by grants from the Landsteiner Stichting voor Bloedtransfusie Research (LSBR-1707 and LSBR-2005),the Dutch Thrombosis Foundation (TSN 2017-01) and the Netherlands Organization for Scientific Research (NWO NWA.1160.18.038). We thank the patients for their contribution and participation in our study.

## Authorship Contributions

MK, IVM, PEB and MH performed research and analyzed data; TK, RF and MF provided vital reagents and expertise; MK, IVM, CM, JV, and RB designed the research and wrote the paper.

## Disclosure of Conflict of Interest

The authors report no conflicts of interest.

## Notes

http://proteomecentral.proteomexchange.org/cgi/GetDataset?ID=PXD027516

### Competing Interest Statement

The authors have declared no competing interest.

### Summary of Updates

Included VWF multimer analysis and targeting of interleukin-8 to Weibel-Palade bodies in NBEAL2 patient-derived endothelial colony forming cells.

