## Supplemental Material for "Mutations in NBEAL2 do not impact Weibel-Palade body biogenesis and Von Willebrand factor secretion in Gray Platelet Syndrome Endothelial Colony Forming Cells"

**Supplemental Materials and Methods**

**Methods**

**Immunoprecipitation and Western blot**

Immunoprecipitation (IP) of GFP and GFP-SEC22B was performed as described.^1^ Western blotting was performed as described ^2^ using primary antibodies for rabbit-anti-NBEAL2 (Abcam [EPR14501], #ab187162 1:1000), mouse-anti-P-selectin (Proteintech, #60322-1-lg, 1:1000), mouse-anti-GFP (Clontech [JL-8] 1:2500), mouse-anti-α-tubulin (Sigma [DM1A] 1:20000), rabbit-anti-VWF (DAKO, #A0085, 1:5000). Secondary antibodies conjugated with infrared dyes (680LT and 800CW) were purchased from LI-COR.

**Immunofluorescence microscopy**

Immunofluorescence (IF) microscopy was performed as described^2^ using the following primary antibodies: rabbit-anti-VWF (DAKO, #A0085, 1:5000), mouse-anti-VWF (CLB [RAg20], described in ^3^, 1:5000), mouse-anti-CD62P-AlexaFluor488 (AbD Serotec [AK-6] 1:100), rabbit-anti-SEC22B (Synaptic Systems, #186003, 1:100), rabbit anti-NBEAL2 (Abcam, [EPR14501], #ab187162, 1:100), rabbit-anti-Rab27A (described in^4^; 1:50), mouse-anti-CD63 (CLB-gran/12, 1:500), goat anti-IL-8 (R&D, #AF-208-NA, 1:150) and goat-anti-Ang2 (R&D, #AF623, 1:200). Secondary antibodies and Hoechst were purchased from Invitrogen (Molecular Probes).

**Secretion assay, ELISA and VWF multimer analysis**

Secretion assays and VWF ELISAs were performed as described.^2,5^ VWF multimer analysis was performed as described previously.^1^ Statistical analyses were performed in Graphpad Prism 8 and are specified in figure legends.

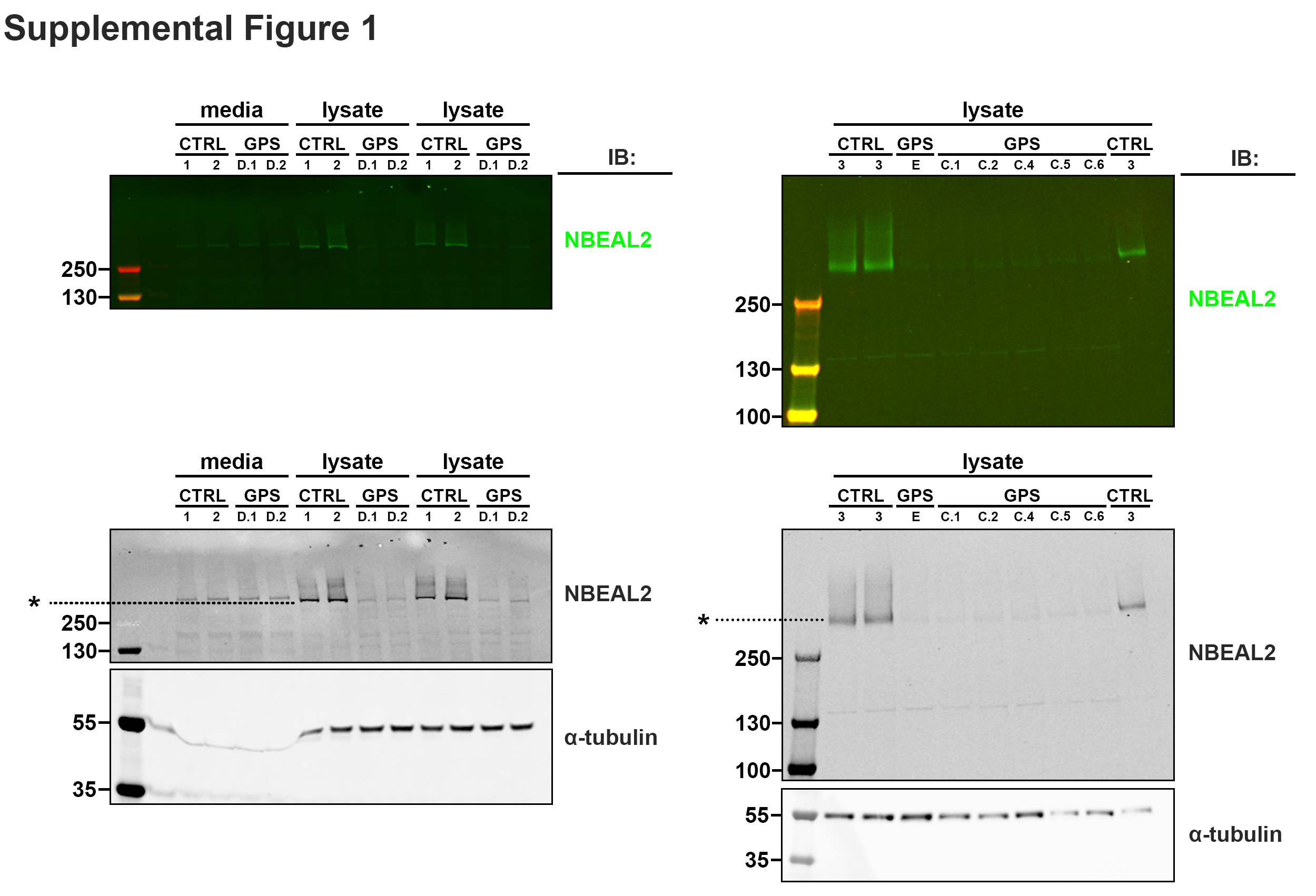

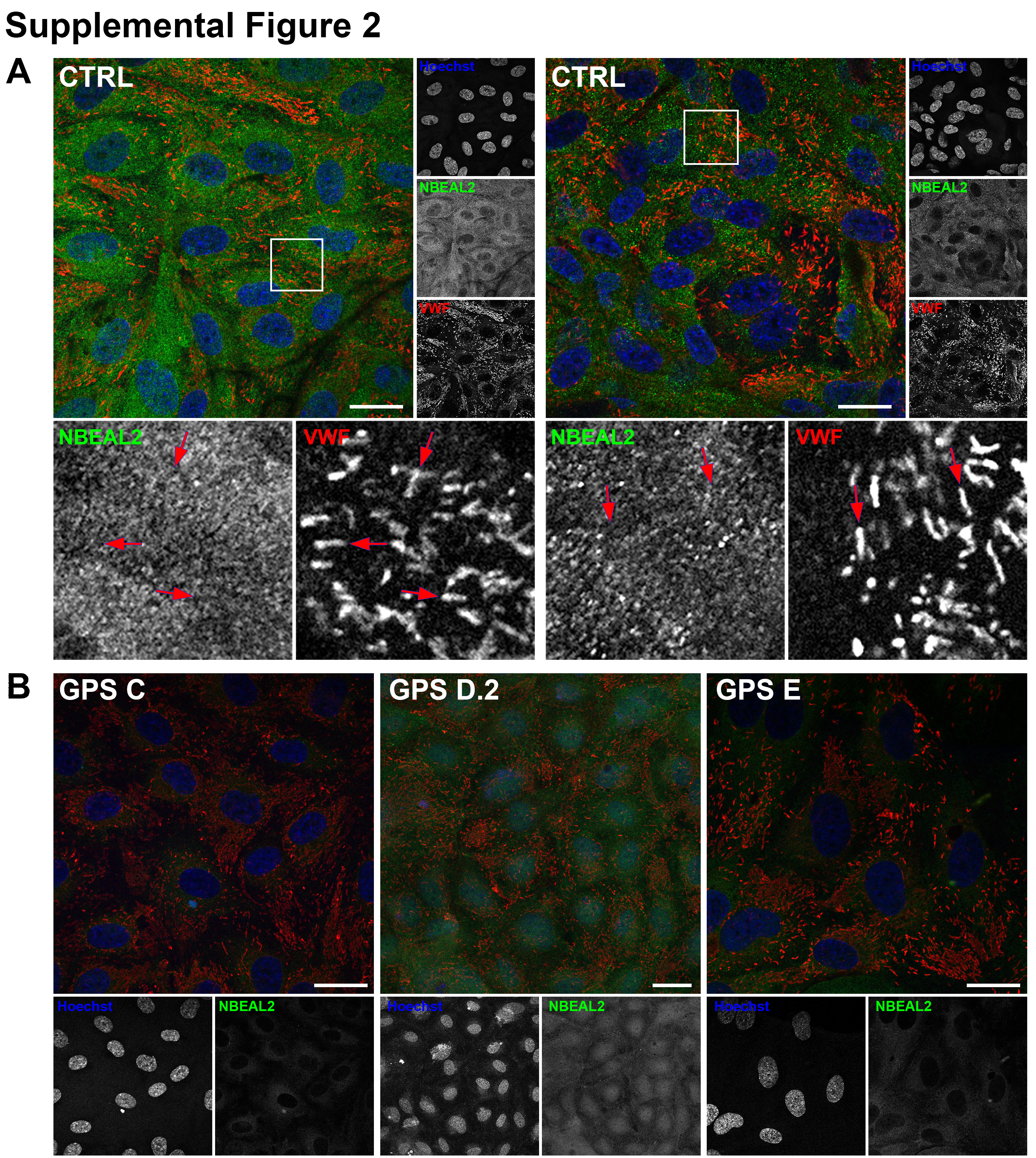

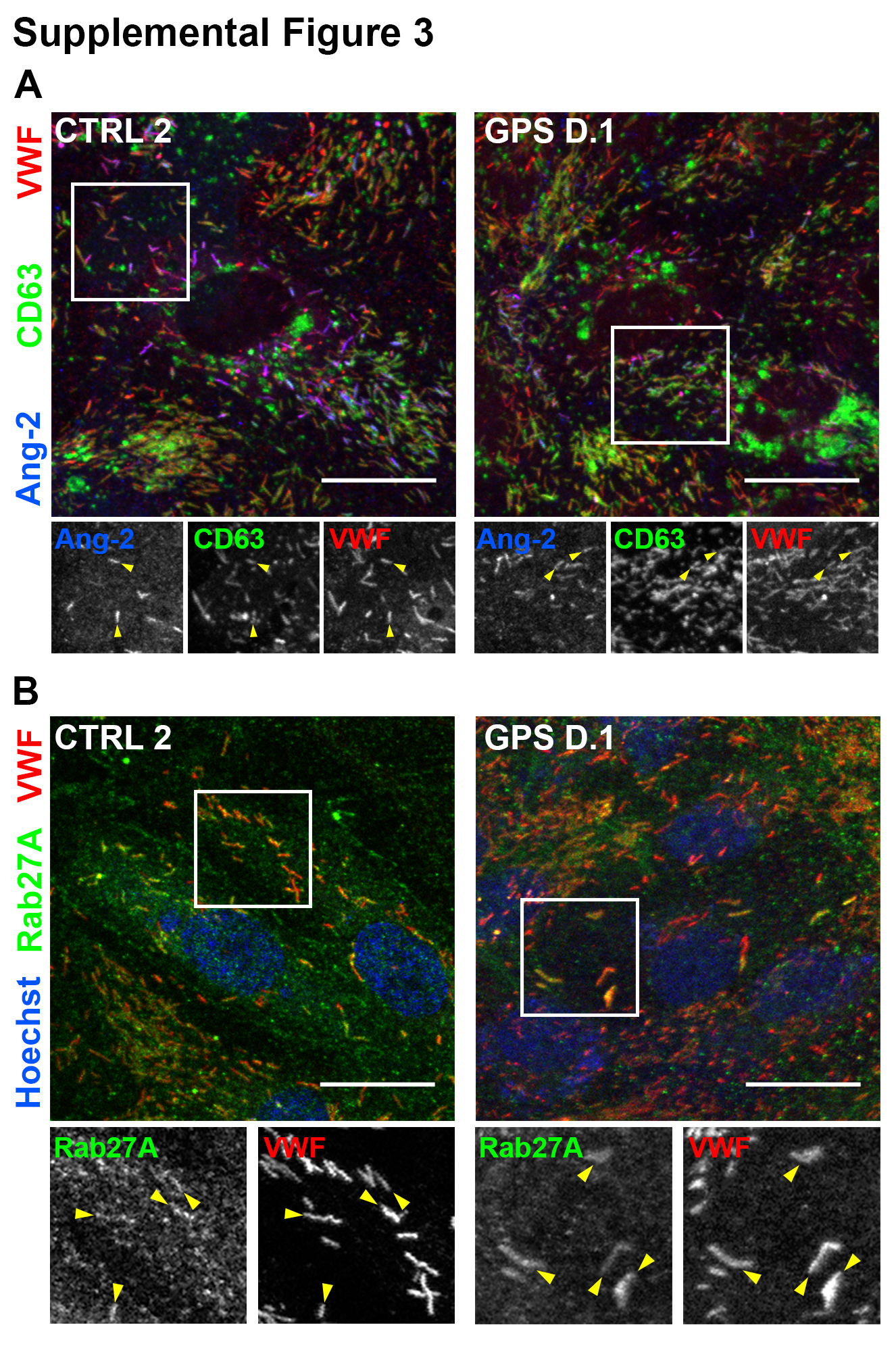

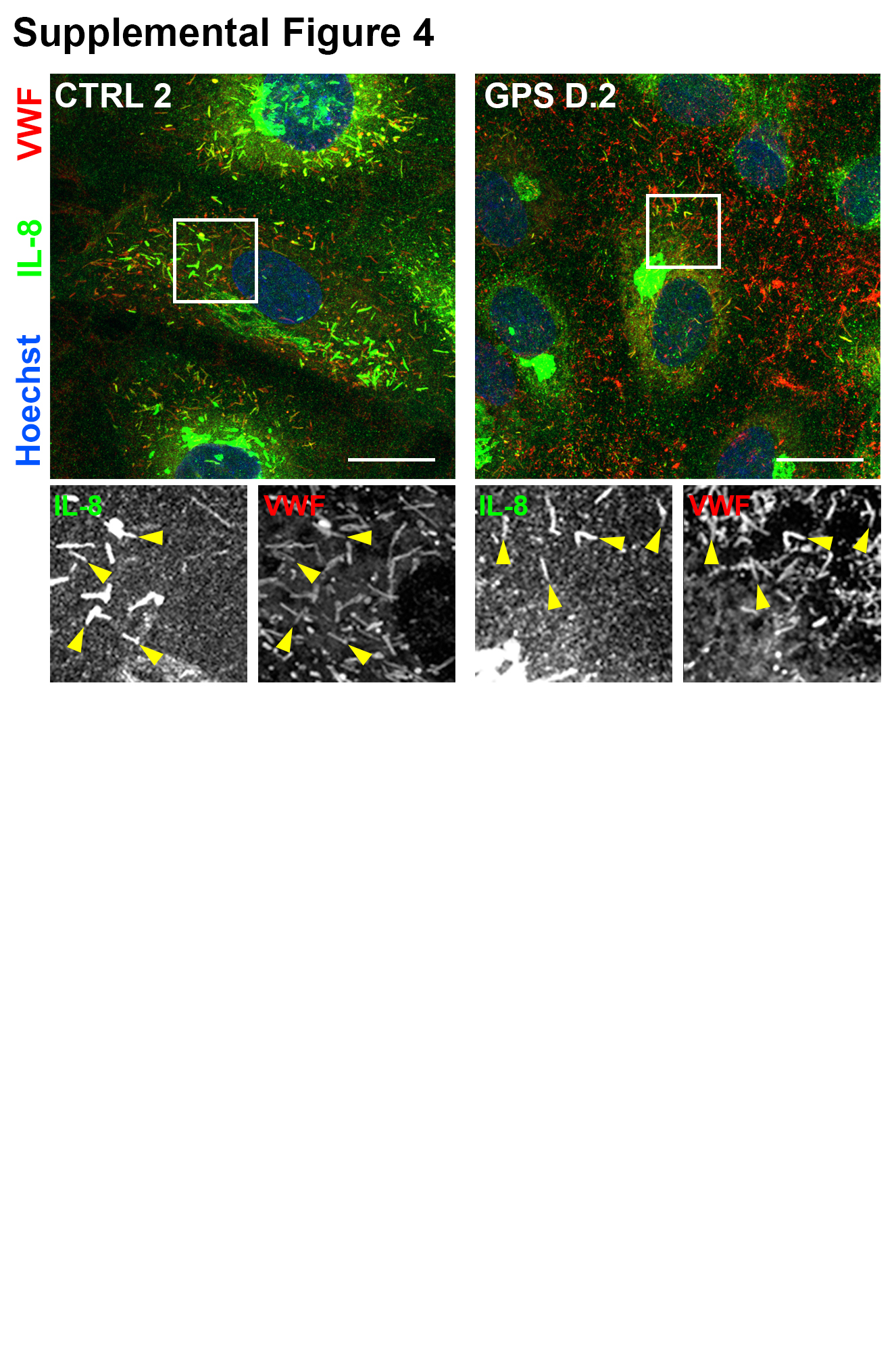

**Supplemental Figure Legends**

**Supplemental Figure 1: NBEAL2 expression in control and GPS ECFCs.**

Western blots of NBEAL2 expression in ECFCs from 3 GPS patients (left: 2 clones (D.1, D.2) were analyzed for patient D; right: 1 clone, E, for patient E and 5 clones for patient C) compared to ECFCs isolated from healthy control donors (CTRL 1-3). α-tubulin is shown as a loading control. Molecular weights of protein ladder marker bands are indicated in kDa on the left. Asterisk indicates full length wild type NBEAL.

**Supplemental Figure 2: NBEAL2 immunostaining in control and GPS ECFCs.**

(A) Representative maximal projections of confocal analysis of control (CTRL) ECFCs, (immuno)stained for VWF (red), nuclei (Hoechst) and NBEAL2 (green; left: polyclonal rabbit anti-NBEAL2, left: monoclonal rabbit anti-NBEAL2). Boxed areas are shown below in grayscale. Scale bars represent 20 µm. Red arrowheads indicate WPBs. (B) Representative maximal projections of confocal analysis of GPS ECFCs of 3 separate GPS patients (C, D and E), (immuno)stained nuclei (Hoechst) and NBEAL2 (green; polyclonal rabbit anti-NBEAL2). Separate channels are shown below in grayscale.

**Supplemental Figure 3: NBEAL2 is dispensable for WPB cargo sorting and maturation.** Representative maximal projections of confocal immunofluorescent analysis of (A) VWF (red), CD63 (green) and Angiopoietin 2 (Ang-2, blue) or (B) VWF (red), Rab27A (green), and nuclei (Hoechst, blue) in CTRL and GPS ECFCs. Boxed areas are shown below in grayscale. Scale bars represent 20 µm. Yellow arrowheads indicate Ang-2 / CD63 positive WPBs.

**Supplemental Figure 4: NBEAL2 is dispensable for inflammatory WPB cargo sorting.** Representative maximal projections of confocal immunofluorescent analysis of (A) VWF (red), interleikin-8 (IL-8, green) and nuclei (Hoechst, blue) in CTRL and GPS ECFCs that were stimulated with 10 ng/ml IL-1β for 20 hours. Boxed areas are shown below in grayscale. Scale bars represent 20 µm. Yellow arrowheads indicate IL-8 positive WPBs.
